# Virus-host interactions on volcanic ash from Mount Etna

**DOI:** 10.64898/2026.06.17.732739

**Authors:** Lisa Voskuhl, Tobias Freitag, Katrin Buder, Mart Krupovic, Sabine Eckhardt, Nikolaos Evangeliou, Luciana Randazzo, Mohamed Tamer Abbara, Sergio Calabrese, Janina Rahlff

## Abstract

Volcanic ash represents an extreme and dynamic habitat, yet it hosts diverse microbial communities with largely unexplored viral diversity. This study investigated bacterial and viral populations in volcanic ash from Mount Etna (Italy) collected during the eruption, focusing on microbial novelty, activity, and virus-host interactions. Taxonomic profiling revealed that *Pseudomonas* and *Telluria* were the dominant bacterial genera, both frequently detected in airborne environments. In contrast, enrichment cultures with volcanic ash were dominated by spore-forming members of the phylum Bacillota, highlighting their resilience under harsh conditions. Metagenomic analysis recovered 19 high-quality metagenome-assembled genomes, including four previously undescribed bacterial species. Replication rate estimates showed that certain taxa were metabolically active, particularly at one sampling site. The presence of Clustered Regularly Interspaced Short Palindromic Repeats (CRISPR) systems with spacers matching viral sequences suggested viral predation pressure on volcanic ash. A total of 1139 viral operational taxonomic units (vOTUs) were identified, of only around half (660 vOTUs) showed similarities to known phages, underscoring the presence of novel viruses. Shared vOTUs across sites revealed the presence of both a core virome and site-specific viral populations. Virus-host predictions indicated frequent interactions with hosts from multiple Gammaproteobacterial genera. Additionally, a 336 kb jumbo phage genome exhibited extensive metabolic capabilities and genetic autonomy. Experimental work identified a unique lytic *Bacillus*-infecting phage (“Phoenix”) with limited propagation capacity. Furthermore, prophage induction experiments revealed active, morphologically diverse temperate phages across multiple bacterial host strains. Overall, these findings highlight volcanic ash as a reservoir of microbial and viral diversity, shaped by environmental extremes and dynamic ecological interactions.

**Highlights:** Metagenomic and cultivation experiments were used to study viruses on volcanic ash

Novelty of viral and bacterial species was detected

Viral-bacterial interactions in metagenomes from volcanic ash were detected

Cultivatable bacteria were mainly spore-forming *Bacilli* species and harbored inducible prophages

## Results & Discussion

### Novelty and diversity of volcanic ash microbes

Volcanoes represent some of the most extreme and dynamic environments on Earth, characterized by high temperatures, steep chemical gradients, toxic acidic gases and volcanogenic particulates, and rapid physical change. Despite these harsh conditions, volcanic systems host diverse and often highly specialized microbial communities ^1,2^. Volcanic lava, on the other hand, has been regarded as mostly sterile ^3^ with microbial life absent and as such newly created volcanic areas can provide insights into microbial colonization ^4^. Far less is known about viruses and microbes inhabiting volcanic emissions. For viruses, most knowledge is derived from hydrothermal environments ^5–7^. Here, we investigate bacteriophages and prophages of volcanic bacterial isolates as well as viruses found in metagenomes sequenced from volcanic ash to characterize viral diversity, host associations, and potential ecological functions in these extreme habitats. Metagenomes from three samples, collected during the fallout at two sites (two samples from site Santa Venerina (SV) and one from Acireale (AC)) around Mount Etna, Sicily, Italy in July and August 2024 were sequenced. The AC site was closer to the coast than the SV site (Figure 1A). Mount Etna is one of the world’s most active open-conduit volcanoes, characterized by continuous passive degassing and frequent explosive eruptions, particularly from the highly active South-East Crater, which produces Strombolian explosions, lava fountains, and widespread ash and lapilli deposits. These basaltic, glass-rich volcanic materials are chemically reactive, enriched in major oxides, nutrients, and potentially toxic trace elements ^8^, and consist of highly heterogeneous, fine-grained particles with vesicular and angular textures (Figure 1B) formed during magma fragmentation and degassing processes. The chemical composition of the ash particles is further described in the Supplement Material. Taxonomic profiling of all ash samples showed a prokaryotic community dominated by Pseudomonadota (66% relative abundance) and Actinomycetota (5.4%) (Figure 1C, Table S1), specifically *Pseudomonas*, *Telluria*, *Sphingomonas aerolata*, and members of the *Acetobacteraceae* were among the most abundant bacterial taxa, ranking within the top 5% of taxa across the three samples (Figure 1D). These bacterial genera are commonly detected in the near-surface atmosphere and on airborne particulate matter ^9,10^ suggesting that they may have been incorporated into the ash plume during atmospheric transport between eruption and deposition. The very different origin of air masses could explain the prokaryotic variability between the three samples (Figure S1). Principal coordinates analysis (PCoA) based on Aitchison distances showed clear separation among the three samples. MDS1 and MDS2 explained 62% and 38% of the variation, respectively, indicating distinct prokaryotic community compositions between the three samples (Figure S2B). By using enrichment cultures, we isolated and sequenced members of the phylum Bacillota, namely those forming clades with *Bacillus subtilis, Lysinibacillus* sp., and *Priestia* sp. (formerly *Bacillus megaterium*) (Figure S3). Most isolates grouped with *Lysinibacillus* spp., while a smaller subset affiliated with *Priestia* spp. and two isolates clustered within the *Bacillus subtilis* clade (Figure S3). *Bacillus* species have been previously detected in volcanic soils ^11^, on lava rocks and in the air of a volcanic island ^12^ and have been shown to increase in abundance on aging lava substrates ^3^. Owing to their ability to form endospores, these bacteria may be particularly well suited for persistence and dispersal in volcanic environments.

**Figure 1:**
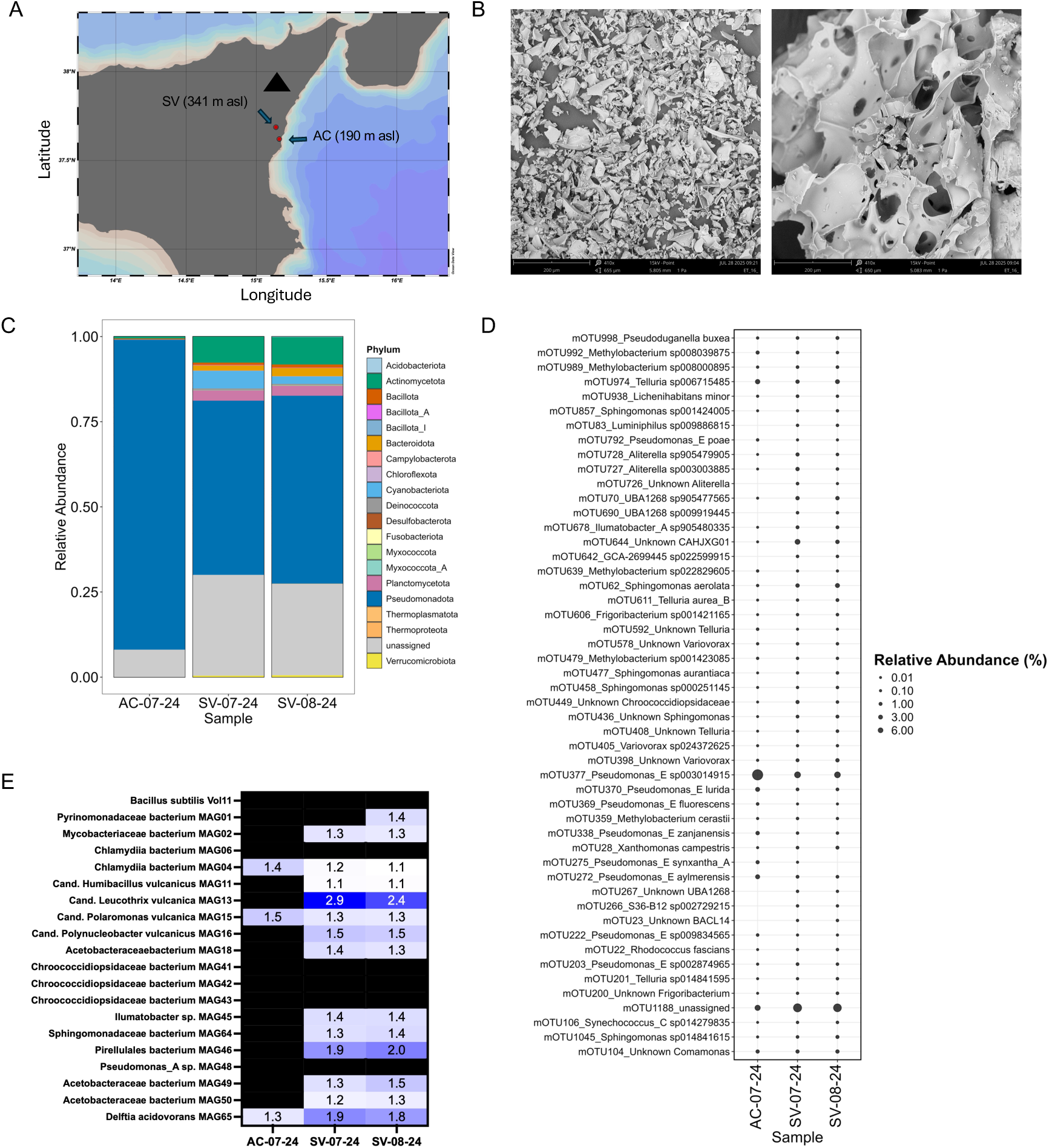
Volcanic ash and sampling sites and prokaryotic community of Etna volcanic ash samples. A, The ash sampling sites on Sicily, Italy close to Mount Etna (black triangle). B, Scanning electron microscopy images of volcanic ash from two different particle-size fractions (0.5-2 mm and <0.074 mm). C, Phylum-level profiling of the prokaryotic community composition of the three ash samples. D, Bubble plot showing top 50 mOTUs and their relative abundance. E, iRep values for metagenome assembled genomes (MAGs) and isolate genome of *Bacillus subtilis* Vol11 indicating actively replicating bacteria.

We recovered 69 metagenome-assembled genomes (MAGs) from the three metagenomic datasets, which were dereplicated to a set of 19 high-quality MAGs (Table S2), four of which represented previously undescribed bacterial species. Based on ANIb values of 80.34 - 83.89% to their closest reference genomes (Table S3), we propose four novel candidate species: Candidatus Humibacillus vulcanicus MAG11, *Cand.* Leucothrix vulcanica MAG13, *Cand*. Polaromonas vulcanica MAG15, and *Cand.* Polynucleobacter vulcanicus MAG16. All MAGs showed high genome quality, with completeness ranging from 94.8 - 99.99% and contamination below 1.3%. The G+C base content of these MAGs varied between 42% and 74% (mean = 57%). Observations of high G+C content has been proposed to correlate with some thermophilic environments, for instance by contributing to increased nucleic acid thermostability ^13^. Using iRep predictions ^14^, we identified the three most actively replicating bacterial populations as *Cand.* Leucothrix vulcanica MAG13 (iRep_max_ = 2.9), *Pirellulales* bacterium MAG46 (iRep_max_ = 2.0), and *Delftia acidovorans* MAG65 (iRep_max_ = 1.9) (Figure 1E). These relatively high iRep values suggest ongoing DNA replication and that these taxa were among the most metabolically active members of the community at the time of sampling. Notably, reliable iRep estimates were obtained predominantly from the two SV sites, implying that microbial populations at these locations were more actively growing or better represented in terms of genome coverage, which is required for robust iRep calculation.

Across all dereplicated MAGs and isolate genomes, we assessed the prevalence of prophages and clustered regularly interspaced short palindromic repeat (CRISPR) systems, which constitute an adaptive immune mechanism in prokaryotes that protects against mobile genetic elements, including viruses ^15^. Both *Cand.* Humibacillus vulcanicus MAG11 and *Delftia acidovorans* MAG65 contained CRISPR arrays with spacers matching viral operational taxonomic units (vOTUs) identified in the metagenomes (Figure 2C, Table S4), demonstrating that these bacterial hosts had previously mounted adaptive immune responses against the corresponding viruses. This observation is consistent with previous findings by Song et al. ^16^, who reported a higher abundance of CRISPR genes in volcanic ash soils compared to forest soils, suggesting that microbial communities in ash environments are exposed to stronger selective pressure from viral predation.

**Figure 2:**
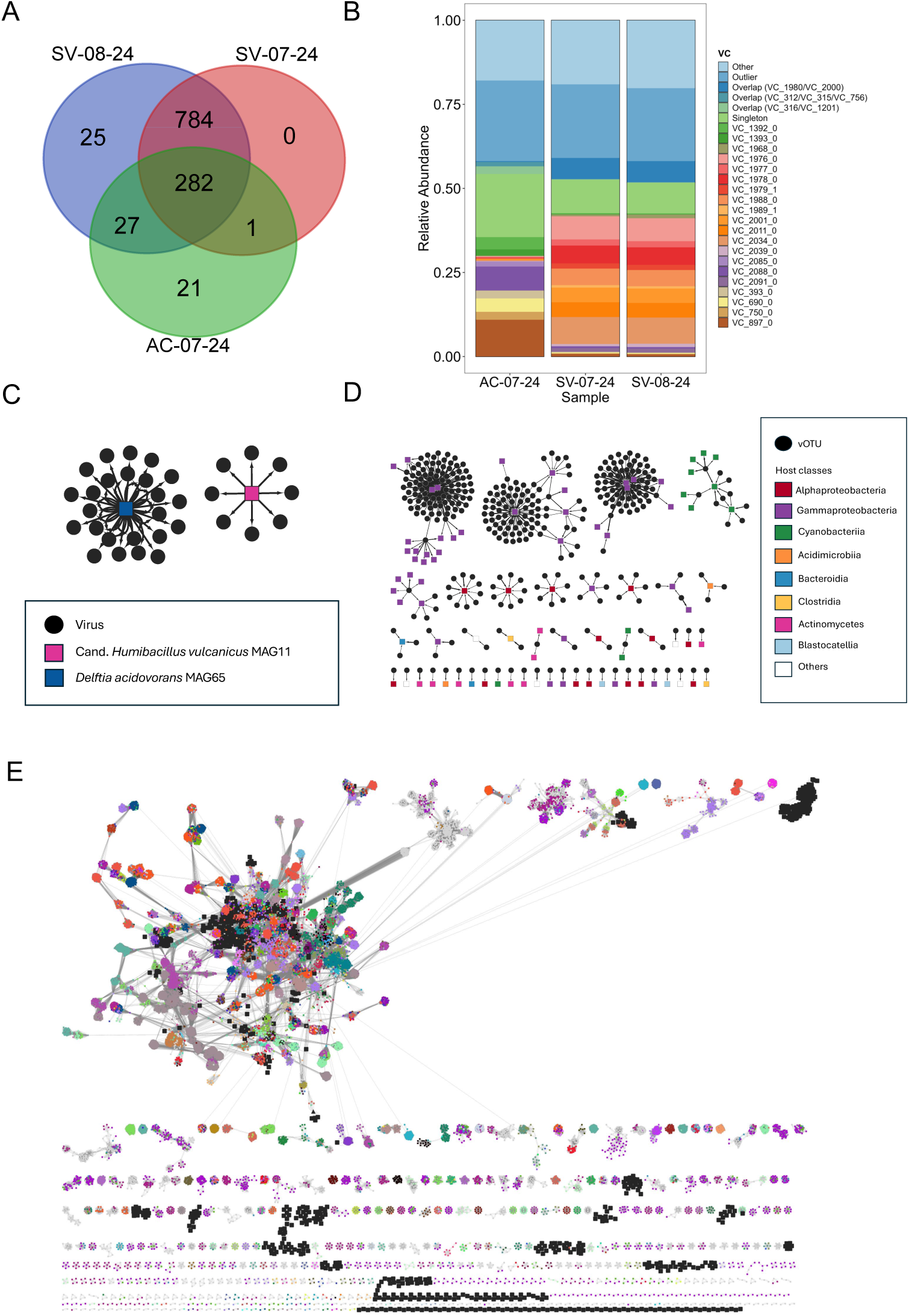
Viral communities and virus-host interactions on volcanic ash samples. A, Venn diagram showing overlapping viral operational taxonomic units (vOTUs) between sampling sites. B, Relative abundance of the top 25 most abundant viral clusters (VC) per sample, the rest assigned as others. C, CRISPR spacer matches from two MAGs to ash-derived vOTUs. D, Virus-host interactions as predicted by iPhoP. E, Protein-sharing network for volcanic ash vOTUs (black squares) clustered with database viruses (colored circles) revealing new VCs and novelty of vOTUs.

### Metagenomic analysis revealed virus-host interactions and a jumbo phage

In total, 1139 vOTUs were detected in the three metagenomes (Table S5), of which 282 were present in all samples. An additional 784 vOTUs were shared between the two SV samples, whereas 25 and 21 vOTUs were site-specific to SV and AC, respectively (Figure 2A). Looking at the relative abundance of vOTUs clustered into genus-level viral clusters (VCs, Table S6) using vConTACT2 revealed clear compositional differences between the two sites. These patterns suggest the presence of a stable core virome on volcanic ash, accompanied by smaller, site-specific viral populations, even though more replicates and samples would be needed to confirm these first trends. Notably, the VC_897_0 cluster, which was abundant at the AC site (11%, only 1% at both SV samples), showed protein similarity to known phages such as Pseudomonas phage DVM-2008 and Halomonas phage phiHAP-1. In contrast, several other abundant clusters (Figure 2B), including VC_1976_0 (7% at SV and 0.3% at AC site), VC_2088_0 (1% at SV and 7% at AC site), and VC_2034_0 (8% at SV and 0.3% at AC site) - displayed no clear similarity to any known phages (Supplementary Table). A distinct viral cluster (the black VC in the upper right corner of Figure 2E), represented a group of previously uncharacterized viruses.

The marked differences in the abundance of viral clusters between the AC and SV sites suggest strong spatial variation in viral community composition. This was further supported by a PCoA analysis showing clear separation among the three samples (MDS1 and MDS2 explained 95.8% and 4.2% of the variation, respectively) indicating distinct viral community compositions between the samples (Figure S2A). The 1139 vOTUs comprised ∼33,000 predicted open reading frames (ORFs). A subset of 692 auxiliary metabolic genes (genes with an AMG flag and a KEGG annotation, Table S7), spanning carbon metabolism, energy generation, nutrient cycling, and redox adaptation suggests that phages actively reprogram host metabolism, as reported before ^17,18^, under the nutrient-limited, metal-rich conditions of volcanic ash. Additional functional capabilities including carbohydrate-active enzymes (277 ORFs, CAZY) and peptidases (544 ORFs) support lytic lifestyles, biofilm degradation, and viral maturation, while recurring toxin-antitoxin (TA) system components such as the HicB_like antitoxin of toxin-antitoxin system (PFAM, <u>PF15970.11</u>) and the Zeta toxin (PFAM, <u>PF06414.18</u>) on 13 and 25 vOTUs, respectively, pointing to roles in persistence and genome stability ^19^ (see Supplementary material for more results and details). Virus encoded TA systems were recently also found on vOTUs from terrestrial steam vents on Hawaii ^20^. Further results on functional annotations of vOTUs can be found in the Supplement material.

In total, 456 virus-host interactions were predicted featuring a confidence score >90 in iPHoP ^21^ (Figure 2D, Table S8). Gammaproteobacteria were the most abundant hosts with the predominant genera being *Pseudomonas* (132 interactions), *Telluria* (69 interactions), and *Comamonas* (62 interactions). In addition, 22 and 43 interactions were predicted with cyanobacteria and Alphaproteobacteria (major genera: *Methylobacterium, Sphingomonas*), respectively. In many cases, the predictions indicated several host genera within the same family for the same virus. While host prediction methods may have limited resolution at the genus level, the frequent assignment of multiple host genera within the same family to individual viruses may also reflect ecological realities of volcanic ash environments, where closely related pioneer taxa dominate and viruses may maintain broader host ranges within host families.

In addition, we detected the genome of a jumbo phage (336,221 bp; 47.1% GC content, 540 open reading frames (Table S9), 100% completeness in CheckV) of which ∼28% (n = 151) could be functionally annotated using sensitive profile-profile comparisons (Figure 3A). The genome was assembled from the August sample of the SV site. Based on read mapping, the jumbo phage genome was also present during July sampling of the same year but not in the coastal AC sample. In the 336 kb jumbo phage genome, multiple structural genes associated with virion assembly, including capsid, tail, baseplate, and tail fiber proteins, are clustered in distinct regions of the genome. The genome encodes a broad repertoire of DNA replication, recombination, and repair functions, such as DNA polymerases (family B and C), helicases (e.g. DnaB-like replicative helicase), recombinases (RecA-like), and small and large subunits of the ribonucleotide reductase, highlighting a high degree of autonomy from host information processing machinery. Additionally, genes involved in nucleotide metabolism (e.g. thymidylate synthase) further support the metabolic independence of this jumbo phage. Notably, several genes typically associated with host-like functions were identified, including tRNAs and transcription-related proteins (e.g., sigma factors), which may enhance translation efficiency and allow better control over host transcriptional processes during infection. The virus also encodes an expanded repertoire of proteins implicated in defense and counter-defense. For instance, it encodes several enzymes involved in metabolism of nicotinamide adenine dinucleotide (NAD), a common target of multiple anti-phage defense systems in bacteria (PMID: 41106366, 39322677). These include nicotinamide/nicotinate mononucleotide adenylyltransferase, glutamine-dependent NAD(+) synthetase, nicotinamide phosphoribosyltransferase, and NADAR-domain N-glycosylase, which could counteract the NAD-dependent ADP-ribosylation-based immune mechanisms. Furthermore, the virus also encodes several endonucleases which could be involved in the degradation of the host chromosome or genomes of coinfecting phages. These results suggest ongoing evolutionary arms-race with the host and possibly competing viruses. The host prediction for this phage was *Hymenobacter* sp., a bacterial genus that was previously found in air ^22^ and surface soil ^23^.

**Figure 3:**
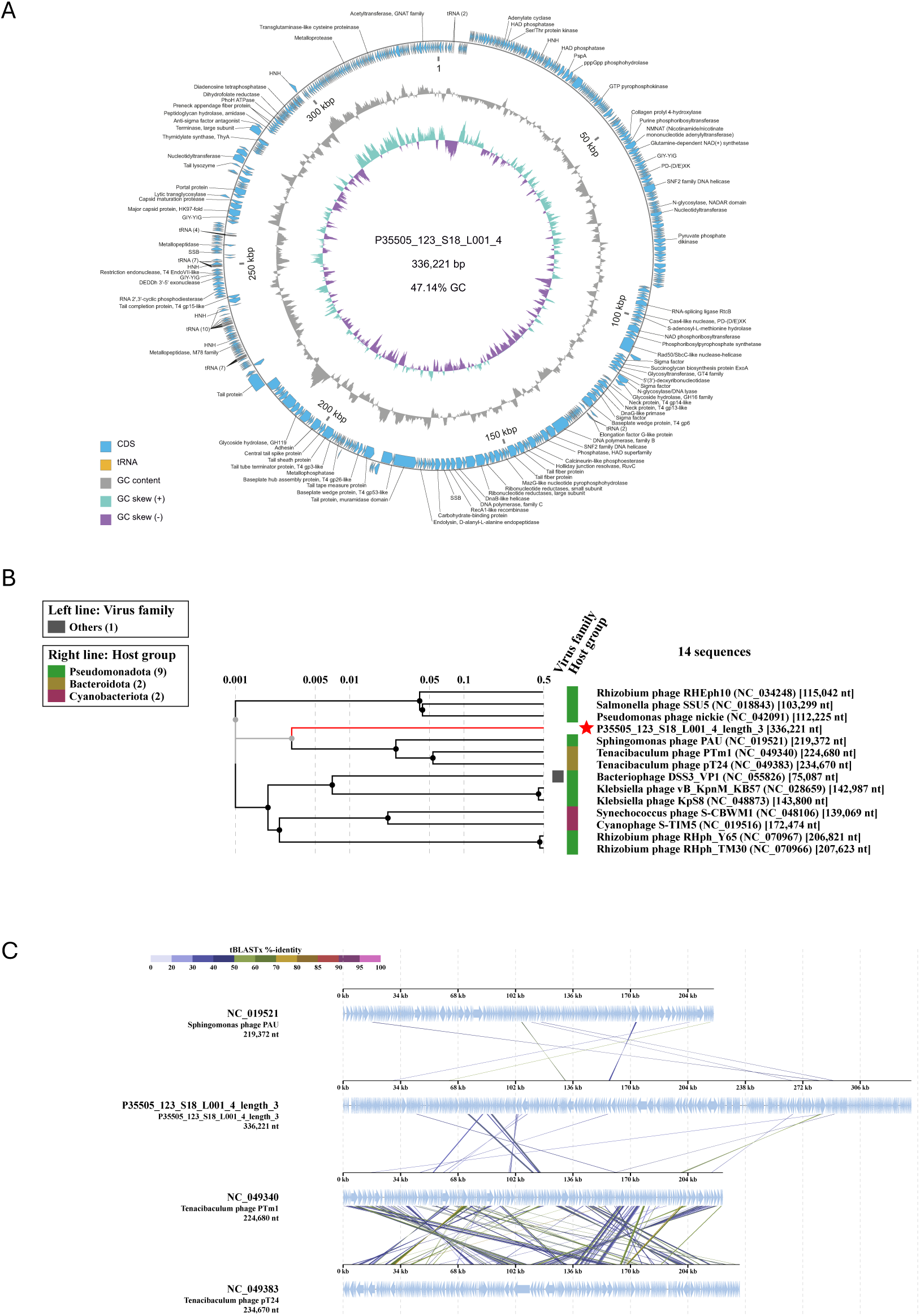
Features and of the jumbo phage genome. A, Map of a 336-kb jumbo phage genome with functionally annotated open reading frames. B, Proteomic tree constructed with ViPTree ^26^ showing placement of the jumbo phage (red star). C, Alignment based on tBLASTx analysis showing identity to known Sphingomonas and Tenacibaculum phages.

The jumbo phage genome had 41% tBLASTx identity over 0.7% length to the known Sphingomonas phage PAU isolated from diseased silkworms (*Bombyx mori* L.) and infecting *Sphingomonas paucimobilis* ^24,25^ (Figure 3A). It had 53.9% tBLASTx identity over 1.9% length to Tenacibaculum phage pT24 and 39% tBLASTx identity over 1.5% length to Tenacibaculum phage PTm1 respectively (Figure 3 B&C). All these phages are also jumbo phages according to their genome sizes (>200 kb). The volcanic jumbo phage shows little similarities to known phages and likely represents a new family. Read coverage to the phage was even (Figure S4), which let us rule out a miss assembly.

### Lytic phage “Phoenix” infecting *Bacillus subtilis* Vol12

We attempted to isolate bacteriophages using 0.2 µm-filtered enrichment cultures derived from volcanic ash (Acireale and Santa Venerina), applying polyethylene glycol (PEG) precipitation and plaque assays with the *Bacillus* isolates as hosts (Table S10). PEG precipitation enabled the visualization of a siphovirus (Figure 4A), whereas plaque assays yielded multiple plaques, many of which were <0.5 mm in diameter. Ultimately, only a single phage was successfully isolated from a plaque on *Bacillus subtilis* Vol12 (isolated on marine agar from ash sample SV from July) with the phage filtrate obtained from the SV sample from August. However, after phage harvesting the phage could not be propagated further on the same host. In transmission electron microscopy (TEM), the phage, designated Bacillus phage Vol12_SV05_R_1 “Phoenix,” displayed a capsid diameter of 44.1 ± 2.5 nm and a tail length of 245 ± 19.1 nm (mean ± SD; *n* = 7 images, Figure 4B & C, Table S11). Its morphology shows myovirus-like sheathed tail (potentially contractile). The slow plaque formation (2-3 days) and the inability to re-propagate “Phoenix” on its host strain pointed to a temperate or slow-replicating lifestyle, a strategy well suited for persistence in the low-biomass, pioneer microbial communities of volcanic ash. Investigating the genome of *Bacillus subtilis* Vol12 further revealed three prophages, which may match the siphovirus morphotypes observed in TEM in the same sample as “Phoenix”, and for which we assume spontaneous induction (further results in supplementary material, Figure S5). Consistently, a number of phages infecting thermophilic variants of *Bacillus* spp. have been described ^27^. While we have not tested if the *Bacillus* strains survive high temperatures or whether heat exposure might facilitate or hinder phage propagation ^28^, it is plausible that the thermal tolerance of the host could influence the induction or replication efficiency of associated phages. Such temperature-dependent dynamics could be particularly relevant in volcanic ash habitats, where environmental conditions fluctuate and may impose additional constraints on phage-host interactions.

**Figure 4:**
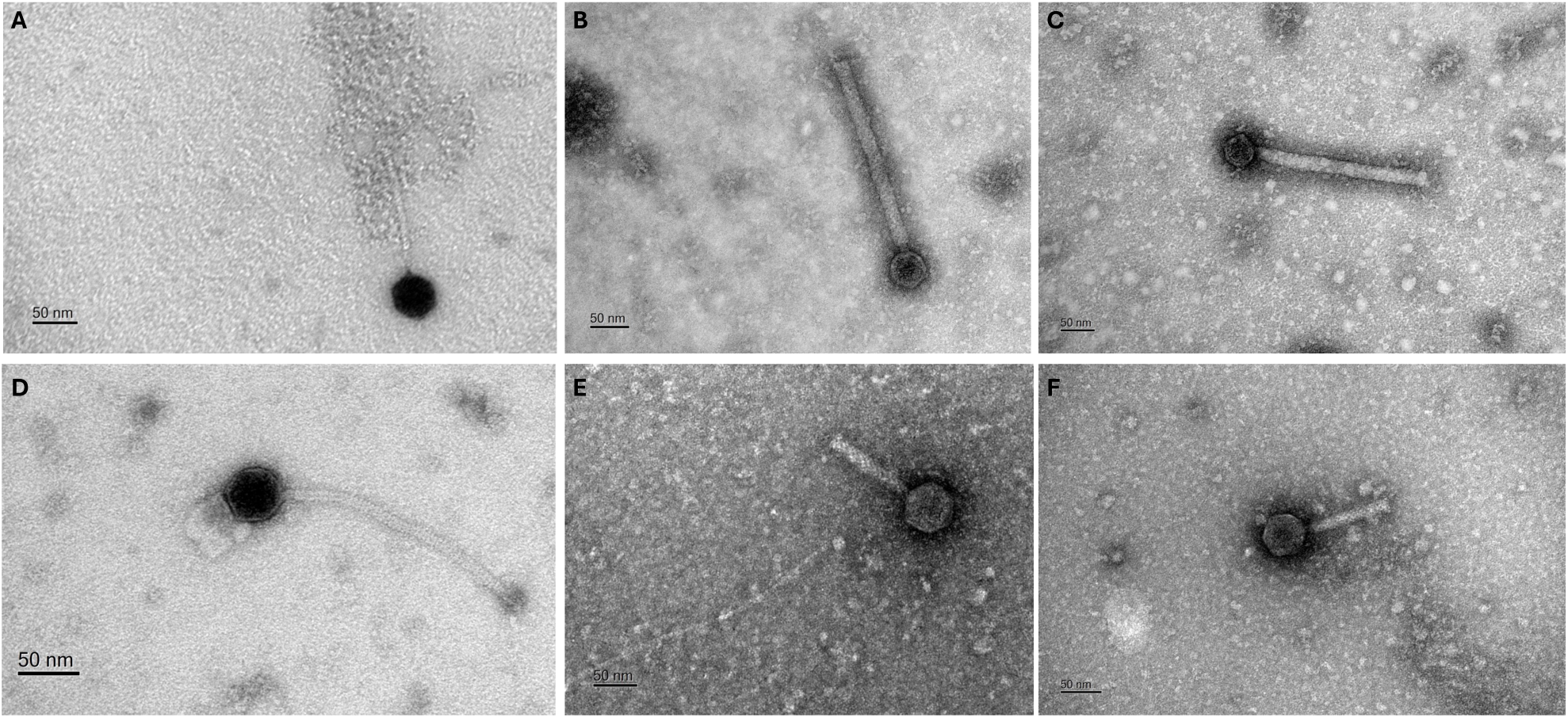
Representative phages observed in TEM. A, Siphovirus after PEG precipitation of ash sample AC inoculated in R2A medium, B+C, *Bacillus* phage Vol12_SV05_R_1 “Phoenix”, and prophages induced from D, *Bacillus subtilis* Vol12, E, *Lysinibacillus* sp. Vol27, and F, *Lysinibacillus* sp. Vol30.

### Morphologically diverse temperate phages in volcanic ash bacteria

Prophages were found in several MAGs: *Cand.* Polaromonas vulcanicus MAG15, *Pseudomonas_A* sp. MAG48, *Delftia acidovorans* MAG65, and *Mycobacteriaceae* bacterium MAG02 and in the genome of *Bacillus subtilis* isolate Vol12. In addition, we chemically induced prophages from representative *Bacillus* strains using mitomycin C (MC) followed by a visual validation using TEM. In successful prophage inductions, growth (as measured by OD_600_) of the strain decreased after ∼4 hours in the presence of high MC concentrations compared to the no MC control. TEM analysis enabled visualization was performed of released viral particles from the isolated strains Vol11, Vol12, Vol18, Vol27, and Vol30 (Figure S6). Prophage inductions in *Bacillus subtilis* Vol11, *Bacillus subtilis* Vol12 and *Priestia* sp. Vol18 yielded siphoviruses (Figure S6, Figure 4D), while *Lysinibacillus* sp. Vol27 and Vol30 yielded myovirus morphotypes (Figure S6, Figure 4E&F), highlighting the presence of morphologically diverse temperate phages in volcanic ash bacterial communities.

In summary, our study reveals that freshly deposited volcanic ash harbored diverse and partially novel microbial and viral communities, despite extremity of this ecosystem. We identified a combination of airborne-derived and ash-adapted bacterial taxa, including several previously undescribed species, alongside metabolically active populations. Viral analyses uncovered a complex viral community consisting of a stable core community and site-specific components, with many viral clusters lacking similarity to known phages, highlighting substantial unexplored diversity. Predicted virus-host interactions and CRISPR evidence indicate ongoing evolutionary dynamics between microbes and viruses in this environment. The detection of a jumbo phage and the isolation of a lytic *Bacillus* phage further demonstrate the functional and ecological complexity of these systems. Additionally, the widespread presence and induction of prophages across multiple endospore-forming hosts of the phylum Bacillota underscore the importance of temperate lifestyles in volcanic ash ecosystems.

### Data and code availability

Metagenomic raw reads and MAGs have been stored in Bioproject PRJNA1364355 at NCBI (Table S12). The 16S rRNA gene sequences are deposited under accessions PZ527938 - PZ527947 in GenBank and at Figshare ^29^. The vOTUs are available under accession numbers PZ033468 - PZ034606 and together with the jumbo phage genome are also deposited at Figshare ^30^. All TEM images are available at Figshare ^31^.

## Acknowledgements

We acknowledge help from Giuseppe Raciti during Etna ash sampling. Data storage was enabled by resources provided by the National Academic Infrastructure for Supercomputing in Sweden (NAISS 2025/22-272 and NAISS 2025/23-14) and the Swedish National Infrastructure for Computing at UPPMAX partially funded by the Swedish Research Council through grant agreement no. 2022-06725. We acknowledge support from the National Genomics Infrastructure in Genomics Production Stockholm funded by Science for Life Laboratory, the Knut and Alice Wallenberg Foundation and the Swedish Research Council, and NAISS/Uppsala Multidisciplinary Center for Advanced Computational Science for assistance with massively parallel sequencing and access to the UPPMAX computational infrastructure. We acknowledge use of the HPC cluster DRACO, with tools and database kindly provided by the VEO group of Bas Dutilh at the FSU Jena. JR received funding by the Swedish Research Council, Starting Grant ID 2023-03310_VR.

## Author contributions

Conceptualization: JR, funding acquisition: JR, methodology: SE, NE, formal analysis: LV, TF, KB, MK, SE, NE, LR, MTA, JR investigation: TF, JR, KB data curation: LV, TF, MK, JR project administration: JR, resources: SC supervision: JR visualization: LV, KB, MK, SE, NE, LR writing – original draft: JR writing – review & editing: all authors

## Declaration of interests

The authors declare no competing interests.

## Star* Methods Sampling

The volcanic ash samples SV-08-24, SV-07-24, and AC-07-24 were sampled from two sites around Mount Etna in Sicily, Italy. They were associated with the paroxysms of Mount Etna that occurred in 2024 between July and August. The sampling sites are in the area close to the volcanic edifice on the eastern slope, and near the municipalities of Santa Venerina and Acireale (approximately 10 and 16 km from the summit craters, respectively). The samples were collected immediately after their deposition on the ground, and each sample therefore refers to a single paroxysmal event. The ash was collected following the IVHHN protocol for the study of leachates^32^. Where possible, samples were collected in a plastic container of known area positioned during ash fall; in other cases, ash was sampled from terraces or roofs, approximately a surface area of one square meter. The ash was collected using powder-free nitrile gloves and a plastic spatula and sealed in plastic bags. At the laboratories of the Dipartimento di Scienze della Terra e del Mare (DiSTeM – University of Palermo), the samples were weighed to obtain the deposition rate (weight/surface area) relative to the paroxysmal event, and then split into different aliquots for morphological, chemical, mineralogical and microbiological analyses.

### Enrichment cultures and 16S rRNA gene sequencing

The three volcanic ash samples (∼200 mg) were added to different types of broth (50 mL per flask), namely Marine Bouillon (Carl Roth, Karlsruhe, Germany), Reasoner’s 2A (R2A, Dinkelberg Analytics GmbH, Gablingen, Germany) and Luria-Bertani (LB) medium placed for up to thirteen days on a shaker at 115 rpm at room temperature. These enrichment cultures were diluted to 10^-2^, 10^-3^, 10^-4^, 10^-5^, and 10^-6^, and up to 500 µL of each dilution was spread onto agar plates containing the respective medium. Approximately 15 mL were filtered through a 0.2 µm syringe filter (Carl Roth syringe filter 0.22 µm, P668.1) and stored in the fridge for use in plaque assay. Plates were monitored for colony formation over several days. Colonies were picked and re-plated thrice for purification. Bacterial genomic DNA from strains Vol11, 12, 18-20, 24-30 was extracted using the E.Z.N.A. ® Tissue DNA kit (Omega Bio-tek, Norcross, USA), and partly concentrated using Zymo Genomic Clean+Concentrator Kit (Zymo Research Europe GmbH, Freiburg im Breisgau, Germany), before quantification using the Qubit dsDNA HS Assay Kit on a Qubit 3.0 Fluorometer (Life Technologies / Thermo Fisher Scientific). A polymerase chain reaction (PCR) was run on the full length 16S rRNA gene using primers 27f (5’-AGAGTTTGATCCTGGCTCAG-3’) and 1492r (5’-TACCTTGTTACGACTT-3’) ordered from Metabion (Planegg, Germany) and by using a Taq PCR Master Mix (2x) (Roboklon, Berlin, Germany). The PCR program was run as follows: Initial denaturation at 95°C, 5 min., 1 cycle; denaturation 95°C, 45 sec., 35 cycles; Annealing at 55°C, 45 sec., 35 cycles; Extension: 72°C, 30 sec., 35 cycles; Final extension: 72°C, 7 min., 1 cycle. The PCR was run on a Biorad Cycler C1000 Touch. The PCR products were checked on an 1.5% agarose gel and purified using DNA Clean+Concentrator TM-K Kit (Zymo Research Europe GmbH) according to the instructions. Sanger sequencing using both the forward and reverse primer was performed at Azenta/GENEWIZ Germany GmbH (Leipzig, Germany). Phylogenetic relationships of volcanic (Vol) isolates were inferred from 16S rRNA gene sequences using the Maximum Likelihood method with the General Time Reversible model. The analysis included 42 sequences (11x Vol isolates, 31x from NCBI) and 1,459 trimmed and aligned positions, with node support assessed by 1,000 bootstrap replicates. The phylogenetic tree was constructed in MEGA12.0.11 ^33^, using Neighbor-Joining and Maximum Parsimony starting trees for heuristic searches and using iTOL v.7 for tree annotation ^34^.

### Whole-genome sequencing of isolate genomes

Whole genome sequencing of the bacterial DNA of selected strains Vol11 and Vol12, was performed at National Genomic Infrastructure (NGI) at SciLifelab in Solna, Sweden using the Illumina DNA method. Samples were sequenced on NovaSeqXPlus (NovaSeqXSeries Control Software 1.3.1.59007) with a 151nt(Read1)-10nt(Index1)-10nt(Index2)-151nt(Read2) setup using ’25B’ mode flowcell. The Bcl to FastQ conversion was performed using bcl2fastq_v2.20.0.422 from the CASAVA software suite. The quality scale used is Sanger / phred33 / Illumina 1.8+. Paired-end Illumina FASTQ files were first interleaved using BBMap’s (https://github.com/BioInfoTools/bbmap) reformat.sh script within bbmap v.39.33.0. Quality control was performed by decompressing the interleaved files and applying adapter trimming and contaminant removal using bbduk.sh from the BBMap suite. Reads were then quality-trimmed and split into paired and singleton outputs using Sickle v.1.33 ^35^ in pe mode and -t sanger setting. The clumpify.sh script within bbmap v.39.33.0.was run to deduplicate the reads. Assemblies were conducted using St. Petersburg genome Assembler (SPAdes) v.3.15.5 ^36^ by using the --isolate flag and filtered to keep only scaffolds >1000 bp.

### DNA extraction from volcanic ash and metagenomic sequencing

DNA was extracted from Mount Etna volcanic ash using PowerSoil Pro Kit (Qiagen, Hilden, Germany). Extractions were performed in triplicates and pooled after the elution step. The concentration was measured in Qubit HS assay test but was below detection limit. Resulting DNA was treated with the Genomic DNA Clean + Concentrator-10 Kit (Zymo Research Europe GmbH) according to the manufacturer’s recommendations and was still beyond the detection limit in Qubit HS assay test. DNA was sent for sequencing with the ThruPLEX DNA-seq method and National Genomic Infrastructure (NGI) at SciLifelab (Solna, Sweden). Samples were sequenced on NovaSeqXPlus (NovaSeqXSeries Control Software 1.3.1.59007) with a 151nt(Read1)-10nt(Index1)-10nt(Index2)-151nt(Read2) setup using ’25B’ mode flow cell. The Bcl to FastQ conversion was performed using bcl2fastq_v2.20.0.422 from the CASAVA software suite. The quality scale used is Sanger / phred33 / Illumina 1.8+.

### Viral metagenomic investigations

Read QC and adapter trimming was performed using bbmap v.39.33 ^37^ under the application of contaminant filtering using the Illumina PhiX spike-in reference genome (phix174_ill.ref.fa) and the artificial contaminants file (sequencing_artifacts.fa). Sickle v.1.33 ^35^ was run in pe mode and -t sanger setting. Since the sequencing approach was PCR-based, the clumpify.sh script (https://github.com/BioInfoTools/BBMap/blob/master/sh/clumpify.sh) within bbmap with setting dedupe subs=1 was used for read deduplication. Scaffolds were assembled for the three samples individually using MEGAHIT v.1.2.9 ^38^, metaSPAdes v.3.15.5 ^39^, and Metaviral SPAdes v.3.15.5^40^. Scaffolds were deduplicated using the dedupe.sh script within bbmap. Finally, the contigs were filtered using pullseq to keep only those >1000 bp. All assemblies per sample were concatenated and seqkit v.2.10.0 ^41^ with rmdup mode was used to remove 100% identical scaffolds. Viral contigs were determined using GeNomad v.1.5.2 ^42^, VirSorter v.2.2.3 ^43^ with option "dsDNAphage,ssDNA,NCLDV,lavidaviridae" and VIBRANT v.1.2.1 ^44^. Resulting viruses were pooled and dereplicated at 95% intergenomic similarity using VIRIDIC v1.0 r3.6. ^45^ Virus-host interactions were predicted using iPhop v.1.4.1 ^21^ using default settings and database version jun_2025_pub. Mapping reads to vOTUs was performed using Bowtie2 v.2.5.4 ^46^ with settings --mp 1,1 --np 1 --rdg 0,1 --rfg 0,1 --score-min L,0,-0.1 for mapping of ≥ 90% identical reads ^47^. Mean coverages of vOTU were determined using the ‘04_01calc_coverage_v3.rb’ script ^48^ and coverage breadth for presence/absence of vOTU (presence at ≥ 75% ^49^) using the calcopo.rb script^50^. Coverages were subsequently normalized to sequencing depth. Relative abundance was expressed as the percentage contribution of each normalized coverage to the overall total. A Venn diagram for vOTUs overlapping between samples was constructed using the UGent tool at https://bioinformatics.psb.ugent.be/webtools/Venn/. Prodigal v.2.6.3 ^51^ in meta mode was applied to predict viral open reading frames. Functional annotations were performed using DRAM-v v.1.5.0 ^52^, and for the jumbo phage genes were functionally annotated using sensitive hidden Markov model profile-profile comparisons with HHsearch v3 (PMID: 31521110). The jumbo phage genome was placed into a proteomic tree and aligned through tBLASTx identity using ViPTree v.4.0 ^26^. The vOTUs were clustered based on shared proteins using vConTACT2 v.0.11.3^53^ using a reference database (release Jan2025) from INfrastructure for a PHAge REference Database (INPHARED) ^54^. Outliers and singleton in vConTACT2 can represent novel viruses. Graphanalyzer.py v.1.6.0 ^55^ was used to compile the virus cluster information.

### Profiling of prokaryotic community and investigating bacterial genomes

Profiling of the prokaryotic community was performed using mOTUs v.4.0.4 ^56^ using the profile option, which maps trimmed metagenomic reads against a the mOTUs database ^57^. Coverages were normalized to sequencing depth by considering average read count and average read length. Relative abundance was calculated as the proportion of each normalized coverage divided by the total, expressed as a percentage. mOTU and vOTU community compositions were analyzed in R using phyloseq v.1.52.0 ^58^. Taxonomic tables, OTU data, and metadata were integrated into a phyloseq object, and relative abundances were calculated on the library-size normalized data. Taxonomic composition was visualized in bar and bubble plots. Beta diversity was assessed using Aitchison distances calculated from compositional abundance data, and patterns in community dissimilarity were visualized by principal coordinate analysis (PCoA). Visualizations were generated using ggplot2 v.3.5.2 ^59^ and microViz v.0.12.7 ^60^.

Binning of metagenome-assembled genomes was performed using MetaBAT2 ^61^ and Maxbin2 v.2.2.7 ^62^ on the assemblies generated with MEGAHIT and metaSPAdes (see above). DAS tool v.1.1.3 ^63^ was used to aggregate the MAGs, which then were manually curated in uBin v.0.9.20 ^48^ and dereplicated using dRep v.3.5.0. in compare and dereplicate mode with default settings ^64^. Quality of MAGs and isolate genomes was checked using CheckM2 v1.0.2 ^65^, and taxonomy was assigned using GTDB-Tk v2.1.1^66^ in classify_wf mode with database version r214. PHASTEST v.1.0.1 ^67^ within Proksee ^68^ was used to detect prophages within MAGs and the isolate genome. CRISPRcasFinder v.1.1.0 ^69^ was used to detected CRISPR systems with evidence level 4 and extract CRISPR spacers from them. CRISPR spacers were matched to ash-derived vOTUs using BLASTn, short algorithm with an 80% similarity filtering step. Mappings were performed on the combined set of dereplicated MAGs and the genome of Vol11 using Bowtie v.2.5.1. ^46^ with --reorder setting. To infer actively replicating bacteria from these mappings, iRep v.1.1.7 ^14^ with setting -mm 3 was subsequently applied. To determine whether the MAGs and isolate genomes correspond to previously undescribed species, we initially identified closely related reference genomes using a tetranucleotide frequency correlation analysis. Subsequently, pairwise average nucleotide identity (ANIb) values were calculated with the JSpeciesWS platform ^70^. Genomes were regarded as putative novel species when their highest ANIb value to any described genome was below 95%. All newly proposed MAG species names have been registered at SeqCode ^71^.

### PEG precipitation from enrichment culture

For polyethylene glycol (PEG) precipitation, cultures in Erlenmeyer flasks containing 50 mL broth (R2A and LB (tryptone 10 g/L, yeast extract 5 g/L, NaCl 10 g/L)) and 200 mg volcanic ash (sample: AC-07-24) were incubated on a shaker at room temperature and 115 rpm for four days (incubation time depending on the culture’s turbidity). Cultures were subsequently filtered through 0.2 µm pore-size filters to remove cells and debris. To 30 mL of the resulting phage-containing filtrate, 7.5 mL of PEG solution (20 g PEG-8000 (Sigma-Aldrich/Merck) and 14.6 g sodium chloride (Carl Roth) dissolved in 100 mL distilled water and sterile-filtered through a 0.2 µm syringe filter ^72^) were added in a 50 mL Falcon tube. The mixture was incubated on ice for 30 min, followed by centrifugation at 10,000 × g for 30 min using a Sorvall RC 6+ Centrifuge (Thermo Scientific). To remove residual PEG solution, last traces of supernatant were gently removed with a micropipette tip. This step was repeated 2-3 times. The pellet containing the phages was gently resuspended in 1000 µL MSM buffer (450 mM NaCl (Carl Roth), 50 mM MgSO_4_ × 7H_2_O (Sigma-Aldrich), 50 mM Trizma base (Sigma-Aldrich) dissolved in 800 mL distilled water, pH adjusted to 8 with HCl, filled to a final volume of 1L with distilled water and autoclaved).

### Plaque assay and phage isolation

The presence of bacteriophages in volcanic ash was investigated by plaque assay using the 0.2 µm filtrate of enrichment cultures of the ash in LB, MB, and R2A broth and the bacterial host strains Vol. 18-20, 24-25, 27-30. For phage detection, 500 µL of each filtrate sample was dispensed onto agar base plates and subsequently covered with 3.5 mL of molten MSM soft agar (450 mM NaCl, 50 mM MgSO_4_ × 7H_2_O, 50 mM Trizma base, 5 g L⁻¹ low-melting agarose) inoculated with 300 µL of the corresponding bacterial culture. After gentle mixing, plates were left to solidify and incubated for 24 - 48 h to allow the development of lysis zones. Putative plaques were excised with sterile pipette tips, transferred to MSM buffer lacking agarose, centrifuged, and stored at 4 °C. For further purification, up to nine streaks per sample were applied to fresh overlay plates containing the appropriate host strain and incubated under the same conditions. Initial screenings showed several clear areas on plates inoculated with different host strains, consistent with possible phage-induced lysis. Plaques were picked with a pipette tip and resuspended in MSM buffer. Phage isolates were subjected to three successive rounds of replating to ensure purification. For phages where this was successful, a serial dilution of the phage against its host was performed, and two plates showing complete lysis were collected and flooded with 5 mL of MSM buffer. The resulting suspensions were clarified by centrifugation at 4500 × g for 20 min., after which the supernatants were passed through sterile 0.2 µm syringe filters and kept at 4 °C. Phage stocks were maintained either as cell-free preparations at 4 °C or as infected host cultures preserved at -80 °C. To generate infected-host stocks, 400 µL of freshly prepared phage suspension was combined with 1.2 mL of an overnight bacterial culture and incubated for 15 min., followed by the addition of glycerol (600 µL 50% glycerol (Sigma) and 900 µL bacterial culture) at -80 °C.

### Prophage induction assays

Prophage induction assays were performed for selected strains (Vol11, Vol12, Vol18, Vol27, Vol30), which were chosen based on taxonomic information and prophage analysis of the genome, if available. Overnight cultures were prepared and aliquoted into 24-well plates (Sarstedt, Nümbrecht, Germany), with 500 µL of culture per well. Prophage induction was initiated by adding a ready-to-use mitomycin C (MC) solution in DMSO (Sigma-Aldrich/Merck, Darmstadt, Germany) at final concentrations of 1, 0.5, and 0.1 µg mL⁻¹, each condition performed in triplicate, alongside untreated controls (no MC). Bacterial growth was monitored by measuring optical density at 600 nm (OD_600_) at approximately hourly intervals in triplicates using a microplate reader Infinite M1000 Pro (Tecan Group Ltd, Männedorf, Switzerland). A pronounced reduction in OD_600_ compared to the control cultures approximately four hours after induction was interpreted as evidence of prophage activation. Following the experiment, cultures showing lysis were pooled and briefly centrifuged. The supernatants were then collected and stored at 4 °C for downstream TEM analysis. For the analysis, each OD_600_ value was normalized by subtracting the initial measurement (t_x_ − t_0_), and all resulting values were plotted against incubation time.

### Transmission electron microscopy of phages

Virus morphology of the lytic phages, the PEG precipitation and the supernatant of the prophage induction were examined by negative-stain ^73^ in TEM. For this purpose, 200-mesh copper grids coated with a Formvar/carbon support film (Science Services GmbH, Munich, Germany) were rendered hydrophilic by glow discharge using a Q150T ES sputter coater (Quorum Technologies, Laughton, UK). A 10 µL aliquot of virus-containing supernatant obtained after prophage induction was placed onto laboratory film (Carl Roth, Karlsruhe, Germany), and the grid was floated film-side down on the droplet. Following adsorption for 1 min, excess liquid was removed by gently touching the grid edge with filter paper. The grid was then immediately transferred onto a drop of 2% (w/v) uranyl acetate solution (pH 4.0-4.5; SERVA Electrophoresis GmbH, Heidelberg, Germany) for 30 sec. After blotting off excess stain, grids were air-dried at room temperature in the dark. Imaging was performed at 80 kV using a JEM-1400 transmission electron microscope (JEOL, Tokyo, Japan) equipped with an Orius SC1000A CCD camera (Gatan Inc., Pleasanton, CA, USA) and operated with Gatan Microscopy Suite software (v2.31.734.0). Capsid and tail dimensions in phages were determined using ImageJ v1.54g in accordance with the measurement approach described by Brum ^74^.

### Scanning electron microscopy (SEM) of ash particles

SEM analyses of ash particles were conducted at the laboratories of the ATeN Centre (Advanced Technologies Network Centre) of the University of Palermo. SEM imaging was used to obtain high-resolution images of the sample surface. A focused electron beam scanned the metallized samples (coated with a thin layer of ultrapure graphite) in a raster pattern, and the signals generated by the beam-surface interaction produced detailed three-dimensional images of surface structures at sub-micrometer resolution (< 1 µm). A Thermo Scientific Phenom XL G2 tabletop scanning electron microscope, equipped with a high-brightness cerium hexaboride (CeB₆) emitter, was used. The instrument operated under both low and high vacuum conditions, with acceleration potentials ranging from 4.8 kV to 20.5 kV. Images were obtained using backscattered electron (BSE) signals, while microanalysis was performed with an energy-dispersive X-ray spectroscopy (EDS) microprobe featuring a silicon drift detector (SDD). The software used for image processing was Alfatest Phenom Pro Suite.

## Notes

### Competing Interest Statement

The authors have declared no competing interest.

