## Supplementary Material for "Virus-host interactions on volcanic ash from Mount Etna"

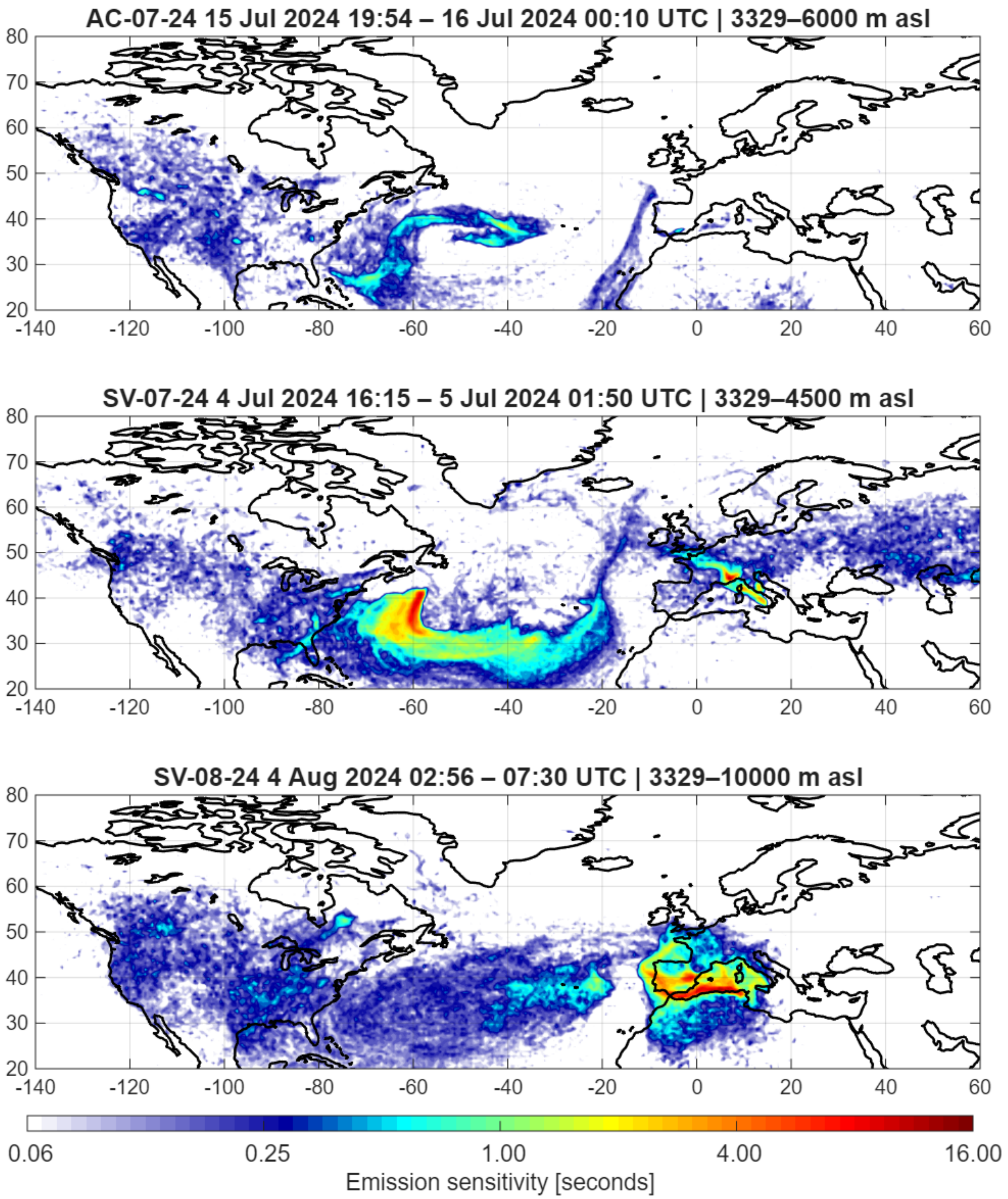


**Figure S1: Backward trajectories for air masses before they reached the Etna.** FLEXPART footprints for the lowest model layer (0-100 m a.s.l.) during the three main phases of the 2024 Etna eruption. Panels show the sensitivity of simulated air masses arriving at Etna for the periods 15-16 July 2024 (top; AC site, release height 3329-6000 m a.s.l.), 4-5 July 2024 (middle; SV site, release height 3329-4500 m a.s.l.), and 4 August 2024 (bottom; SV site, release height 3329-10000 m a.s.l.). Footprints are integrated over the indicated eruption periods and release heights. Colors indicate emission sensitivity in seconds.

### Methodology for Etna footprints

Source-receptor sensitivities (so-called "footprints") were calculated using the Lagrangian particle dispersion model FLEXPART version 11 [1] in backward mode from the receptor location (Etna, 3329 m asl). FLEXPART traces particles backward in time from the receptor and yields a gridded footprint, which describes the influence of upwind surface emissions on the sampled air mass at the receptor. For the Etna eruptions in 2024, 50000 particles were released at 3329-4500 m a.s.l., as 4500 m a.s.l. was the estimated top-height of the plume between 4/7/2024 at 16:15 and 5/7/2024 at 01:50. Accordingly, the same number of particles was released at 3329-6000 m a.s.l between 15/7/2024 at 19:54 and 16/7/2024 at 00:10, and at 3329-10000 m a.s.l. on 4/8/2024 between 02:56 and 07:30. FLEXPART was driven by operational hourly, 0.5° horizontal resolution meteorological fields from the ERA5 reanalysis [2]. It considers several processes along the trajectories, such as turbulence [3], unresolved mesoscale motions [4], and convection [5].

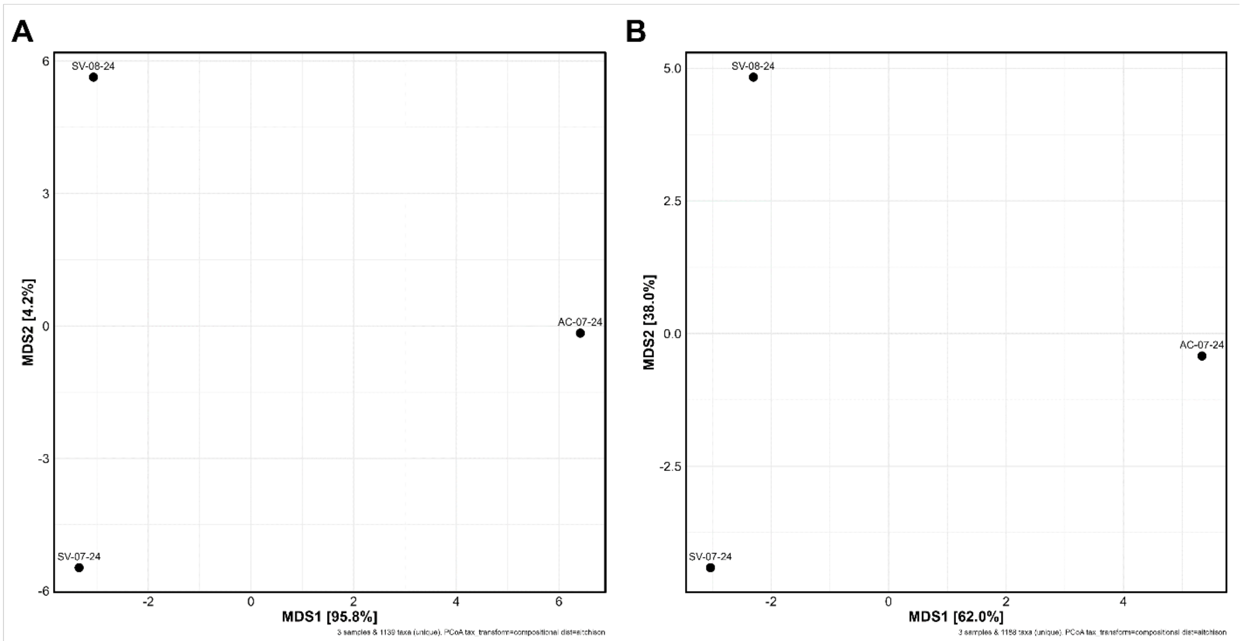


**Figure S2: Principal coordinates analysis (PCoA) showing differences in community composition among samples from one AC and two SV sites based on Aitchison distances.** A, Viral community composition (vOTUs), showing clear separation among the three samples, with

MDS1 explaining 95.8% of the variation and MDS2 explaining 4.2%, indicating strong spatial variation in viral cluster abundance between sites. B, Prokaryotic community composition (mOTUs), also showing distinct separation among samples, with MDS1 explaining 62% of the variation and MDS2 explaining 38%, indicating marked differences in microbial community composition between the AC and SV sites.

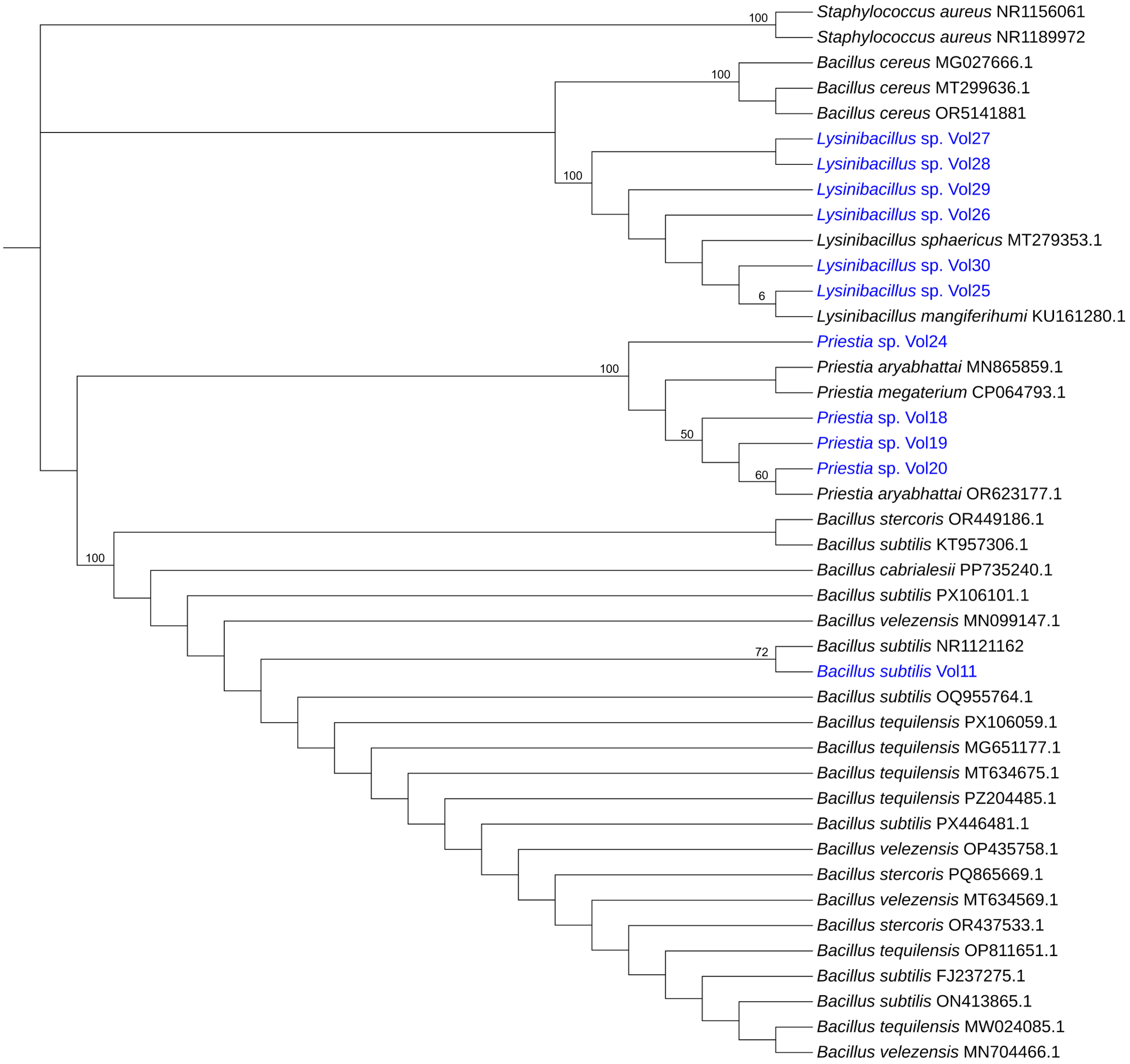


**Figure S3: Phylogenetic tree for 16S rRNA genes from Vol isolates.** Phylogenetic tree inferred using the Maximum Likelihood method based on 42 nucleotide sequences with 1,459 aligned positions. Bootstrap support values (%) are shown at the nodes based on 1,000 replicates, with branches supported by less than 50% collapsed. The initial tree for heuristic searching was selected based on the highest log-likelihood from Neighbor-Joining and Maximum Parsimony starting trees. Evolutionary distances were computed using the General Time Reversible model. Analyses were performed in MEGA12 [6].

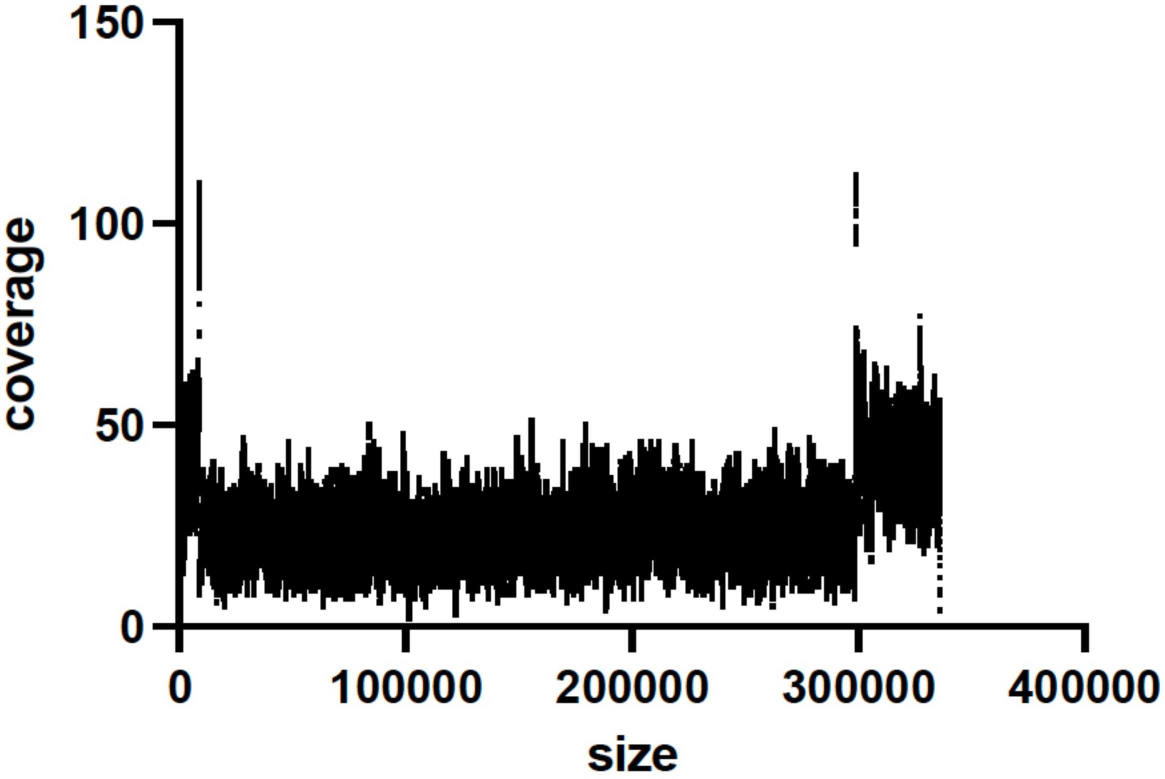


**Fig S4: Coverage profile for jumbo phage genome.** Reads were mapped to the genome using Bowtie2 [7].

**Mount Etna and chemical composition of volcanic ash**

Mount Etna is one of the most active open-conduit volcanoes in the world, continuously emitting large amounts of acidic gases and potentially toxic elements (PTEs) into the atmosphere [8]. Passive degassing is frequently alternated with explosive and effusive activity from the four summit craters: Voragine, Bocca Nuova, North-East Crater, and South-East Crater. Over the past two decades, the South-East Crater has been the most active, characterized by intense explosive activity including Strombolian explosions, lava fountains, and paroxysmal events. During explosive episodes, large quantities of volcanic ash and lapilli are emitted and deposited across both local and regional scales. These materials are predominantly basaltic and glass-rich, with high surface area, vesicular porosity, and a chemically reactive matrix containing major oxides and trace nutrients. The principal components include $SiO_2$ (45-52%), $Al_2O_3$ (13-18%), FeO/$Fe_2O_3$ (8-12%), CaO (8-12%), MgO (5-10%), and alkalis ($Na_2O$ + $K_2O$, 3-6%). Minor and trace elements, often associated with water-soluble surface salts (mainly sulfates, chlorides, and fluorides), include P, Mn, Zn, Cu, Ni, and significant concentrations of PTEs (e.g., As, Ba, Bi, Cd, Cr, Cs, Mo, Pb, Sb, Se, Te, Th, Tl, U, V). Ash and lapilli morphology is highly heterogeneous, consisting mainly of angular volcanic glass, crystals, and lithic particles. Irregular particle surfaces and vesicular textures are produced during magma degassing and fragmentation processes. From a granulometric perspective, ash falls within the <2 mm fraction but is typically dominated by fine particles (micron- and submicron-sized), with a broad size distribution reflecting fragmentation and transport processes.

**Further results on DRAMv functional annotations**

DRAMv functional annotations of the full set of vOTUs from Mount Etna volcanic ash metagenomes yielded 6,876 functionally annotated ORFs matching VOGDB entries, the majority were classified as unknown function (Xu; n=3,825), followed by hallmark viral proteins (Xh; n=2,307), replication-associated (Xr; n=1,186), structural (Xs; n=1,099), and packaging functions (Xp; n=538). Additionally, 1,913 ORFs were flagged as VOGDB-annotated (V). AMG flag analysis identified 692 AMG candidates carrying an AMG flag and a KEGG annotation, containing 675 unique KEGG hits, 38 unique CAZyme hits, and 2137 unique PFAM hits. These AMGs suggest that phages actively reprogram host metabolism to improve replication efficiency under the nutrient-limited, metal-rich conditions of volcanic ash. Functionally annotated genes involved in carbon and central metabolism (e.g. glyceraldehyde-3-phosphate dehydrogenase, enolase, malate dehydrogenase, fructose-1,6-bisphosphatase, transaldolase) and energy generation (e.g. NADH dehydrogenase, cytochrome c oxidase, succinate dehydrogenase) point to enhanced host energy and carbon flux. Further AMGs spanning sulfur and nitrogen cycling, redox and stress adaptation (e.g. arsenite methyltransferase, nitric oxide reductases, superoxide dismutase, glutathione-related enzymes), and cofactor/vitamin biosynthesis (e.g. biotin carboxylase, cobalamin biosynthesis protein) indicate broad viral capacity to optimize host survival in a geochemically harsh environment. MFS-type multidrug resistance transporters may enhance host tolerance to the toxic metal characteristics of volcanic substrates, while 250 ORFs carrying heme regulatory motifs suggest viral interactions with iron acquisition pathways, which might be particularly relevant given the iron-rich volcanic substrate.

The vOTUs displayed clear modular genome organization, combining hallmark structural and replication genes, including portal proteins, major capsid proteins, tail fiber components, endolysins, and spanin-like proteins, with regulatory and stress-associated elements. Thirteen scaffolds contained genes for the HicA/HicB toxin–antitoxin (TA) system, where HicA functions as an mRNA interferase inhibiting translation via RNA cleavage and HicB serves as its cognate antitoxin [9]. Across five additional scaffolds, a Zeta toxin paired with an AAA ATPase domain was identified, which is associated with plasmid maintenance and bacterial persistence in biofilm formation. Together, these TA systems likely contribute to phage persistence, genome maintenance, and stabilization of mobile genetic elements under environmental stress [9]. Complementing these findings, LuxR-family transcriptional regulators suggest quorum-sensing-mediated coordination of gene expression linked to stress-response pathways [10,11], SOS-associated peptidases addressing DNA damage response systems [12], and deglycase-like enzymes involved in metabolic repair processes [13]. Protein similarity searches revealed that the viral community spans both bacteriophage and eukaryotic virus lineages. Similarities of functionally annotated genes to those of giant viruses such as *Fadolivirus algeromassiliense* (Mimivirales), bacteriophages infecting *Ralstonia*, *Burkholderia*, *Pseudomonas*, and *Halomonas*, and protist-associated viruses such as *Bodo saltans* virus and *Pyramimonas orientalis* virus, suggesting that microbial eukaryotes may also play a role in early colonization processes of volcanic ash.

Overall, volcanic ash from Mount Etna hosts a highly diverse, functionally rich, and largely novel viral community. These phages encode a broad repertoire of metabolic, structural, and regulatory genes likely to shape host physiology and ecosystem processes under extreme conditions, positioning volcanic deposits as underexplored hotspots of viral diversity and evolution.

**Results on prophages of *Bacillus subtilis* Vol12**

*Bacillus subtilis* Vol12, the host of the lytic phage Vol12_SV05_R_1 "Phoenix," additionally contained three prophages in its genome. Prophage(s) from Vol12 could be induced with Mitomycin C (see main text and results below). In addition, during plating of the host strain *Bacillus subtilis* Vol12, few occasional plaques appeared (Figure S5A). Spontaneously appearing plaques in seeded Vol12 plates, when picked and used in plaque assay against the host strain, do not show up again. In the harvested supernatant containing the lytic Phoenix phage, which has a myovirus morphotype, few additional siphoviruses were observed in transmission electron microscopy (Figure S5 B+C), supporting the spontaneous induction of prophage(s).

Prophage sequences within the host genome were detected using PHASTEST v.1.0.1 [14] with Deep/PHAST-BSD annotation mode within Proksee webserver [15], and further investigated using the Viral Proteomic Tree server (ViPTree) v.4.0 [16] with database settings host type: Prokaryote, nucleotide type: dsDNA. We found that Prophage 1 and 2 mainly shared protein identities with each other (70.1% mean identity in tBLASTx over 52.8% length), whereas Prophage 3 shared proteins with known Bacillus phage phi105 (accession NC_048631, 68.7% mean identity over 57.3% length) (Figure S2 D+E). All three prophages were placed within clusters of existing *Bacillus* prophages.

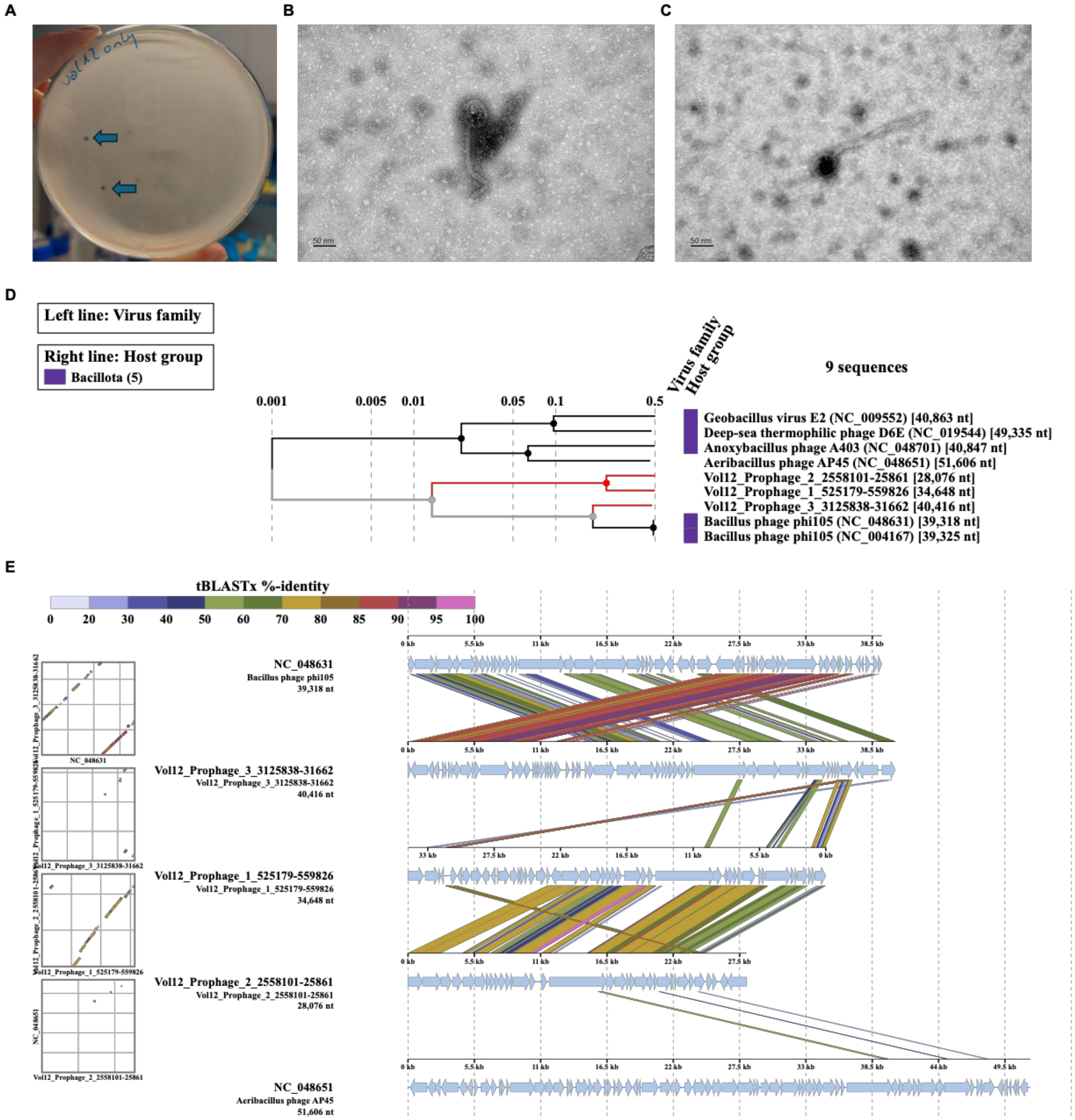


**Figure S5: Prophages of *Bacillus subtilis* Vol12.** A, Two plaques in host strain *Bacillus subtilis* Vol12 indicated by blue arrows. B+C, Siphovirus morphotypes in the harvested supernatant for Vol12_SV05_R_1 “Phoenix”. D, Proteomic tree for clustering of Vol12 prophages based on tBlastX comparison method. E, Protein-based alignment of Vol12 prophages based on tBlastX comparison method with each other and two known related phages.

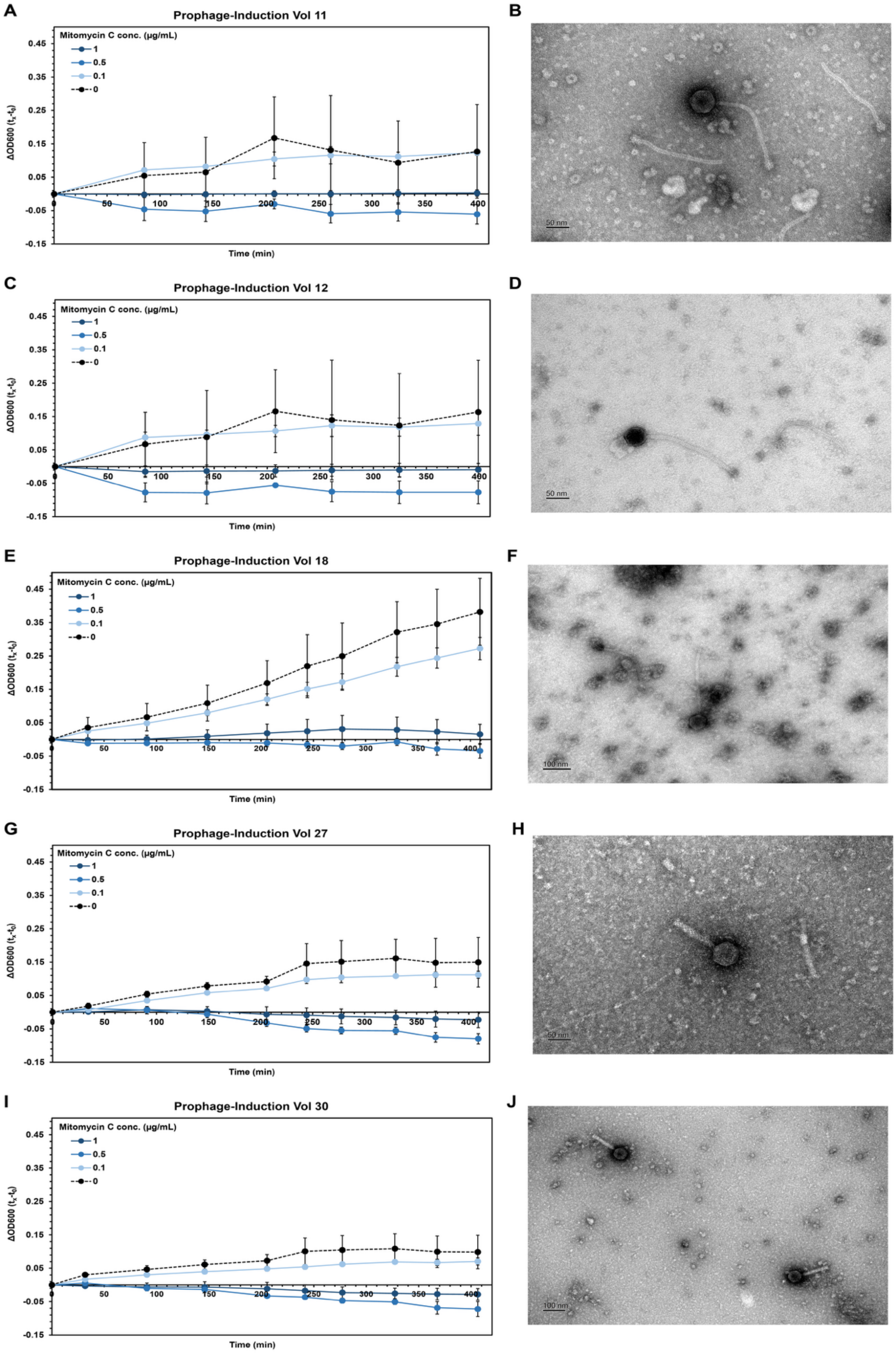
A
Prophage-Induction Vol 11
Mitomycin C conc. (µg/mL)
1
0.5
0.1
0
ΔOD600 (tx-t0)
Time (min)
B
50 nm
C
Prophage-Induction Vol 12
Mitomycin C conc. (µg/mL)
1
0.5
0.1
0
ΔOD600 (tx-t0)
Time (min)
D
50 nm
E
Prophage-Induction Vol 18
Mitomycin C conc. (µg/mL)
1
0.5
0.1
0
ΔOD600 (tx-t0)
Time (min)
F
100 nm
G
Prophage-Induction Vol 27
Mitomycin C conc. (µg/mL)
1
0.5
0.1
0
ΔOD600 (tx-t0)
Time (min)
H
50 nm
I
Prophage-Induction Vol 30
Mitomycin C conc. (µg/mL)
1
0.5
0.1
0
ΔOD600 (tx-t0)
Time (min)
J
100 nm

**Figure S6: Prophage induction experiments and TEM images of induced phages** for A, strain *Bacillus subtilis* Vol11, B, induced siphovirus for *Bacillus subtilis* Vol11, for C, strain *Bacillus subtilis* Vol12, D, induced siphovirus for *Bacillus subtilis* Vol12, E, for *Priestia* sp. Vol18, F, induced siphovirus for *Priestia* sp. Vol18, G, strain *Lysinibacillus* sp. Vol27, H, induced siphovirus for *Lysinibacillus* sp. Vol27, I, strain *Lysinibacillus* sp. Vol30, J, induced siphovirus for *Lysinibacillus* sp. Vol30.
